# A cell-free platform for rapid synthesis and testing of active oligosaccharyltransferases

**DOI:** 10.1101/145227

**Authors:** Jennifer A. Schoborg, Jasmine Hershewe, Jessica C. Stark, Weston Kightlinger, James E. Kath, Thapakorn Jaroentomeechai, Aravind Natarajan, Matthew P. DeLisa, Michael C. Jewett

## Abstract

Protein glycosylation, or the attachment of sugar moieties (glycans) to proteins, is important for protein stability, activity, and immunogenicity. However, understanding the roles and regulations of site-specific glycosylation events remains a significant challenge due to several technological limitations. These limitations include a lack of available tools for biochemical characterization of enzymes involved in glycosylation. A particular challenge is the synthesis of oligosaccharyltransferases (OSTs), which catalyze the attachment of glycans to specific amino acid residues in target proteins. The difficulty arises from the fact that canonical OSTs are large (>70 kDa) and possess multiple transmembrane helices, making them difficult to overexpress in living cells. Here, we address this challenge by establishing a bacterial cell-free protein synthesis platform that enables rapid production of a variety of OSTs in their active conformations. Specifically, by using lipid nanodiscs as cellular membrane mimics, we obtained yields of up to 440 µg/mL for the single-subunit OST enzyme, ‘Protein glycosylation B’ (PglB) from *Campylobacter jejuni*, as well as for three additional PglB homologs from *Campylobacter coli, Campylobacter lari*, and *Desulfovibrio gigas*. Importantly, all of these enzymes catalyzed *N*-glycosylation reactions *in vitro* with no purification or processing needed. Furthermore, we demonstrate the ability of cell-free synthesized OSTs to glycosylate multiple target proteins with varying *N*-glycosylation acceptor sequons. We anticipate that this broadly applicable production method will advance glycoengineering efforts by enabling preparative expression of membrane-embedded OSTs from all kingdoms of life.

## Introduction

Asparagine-linked (*N*-linked) glycosylation is one of the most prevalent polypeptide modifications in nature and is present in all domains of life (Apweiler, Hermjakob, & Sharon, 1999; Nothaft & Szymanski, 2010). Glycosylation affects a multitude of protein characteristics including folding, immunogenicity, activity, half-life, and regulation of signaling cascades (Apweiler et al., 1999; Helenius & Aebi, 2004; Varki, 1993; Walsh, 2010). Despite the importance of glycans in biology, defining the rules governing structural and functional consequences of site-specific glycosylation remains an active area of investigation. Challenges arise due to several complications associated with producing and characterizing natural glycosylation systems, however. One of the major challenges is that methods for synthesis and functional analysis of enzymes that modify and transfer glycans remain limiting, as exemplified by the small fraction of characterized enzymes (1% of more than 250,000) in the carbohydrate-active enzymes database (http://www.cazy.org/) (Lombard, Ramulu Golaconda, Drula, Coutinho, & Henrissat, 2014).

Oligosaccharyltransferases (OSTs) are a class of membrane-bound enzymes that contain many (up to 13) transmembrane helices. OSTs facilitate the transfer of preformed oligosaccharides *en bloc* from lipid-linked oligosaccharides (LLOs) to target proteins bearing an amino acid sequence, or sequon, of the form N-X-S/T, where X is any amino acid except proline. In higher eukaryotes, the OST is composed of multiple subunits with the catalytic subunit denoted as STT3, whereas known archaeal and bacterial OSTs are composed of a single subunit, typically denoted as AglB and PglB, respectively (Kelleher & Gilmore, 2006; Maita, Nyirenda, Igura, Kamishikiryo, & Kohda, 2010; Matsumoto et al., 2013). OSTs are effectively the ‘‘gatekeepers” of *N*-linked glycosylation because their substrate preferences determine *which* proteins will be glycosylated, *where* the glycosidic bond will be formed, and *which* oligosaccharide will be attached at a given site (Chen, Glover, & Imperiali, 2007; Kowarik, Young, et al., 2006; Lizak, Gerber, Numao, Aebi, & Locher, 2011; Ollis et al., 2015). Therefore, the ability to rapidly synthesize and characterize a wide range of OSTs would be a valuable biochemical tool that could enable a deeper understanding of these important enzymes and unlock their full biotechnological potential. Unfortunately, recombinant expression of integral membrane proteins in living cells is tedious, and is limited by issues arising from cell toxicity and insolubility (Jaffee & Imperiali, 2013). Furthermore, purification and refolding of membrane proteins often requires lengthy optimization. These challenges present a unique opportunity for the application of cell-free protein synthesis (CFPS) in combination with *in vitro* glycosylation (IVG) systems for preparative expression and functional characterization of OSTs.

CFPS in crude lysates provides an alternative method for producing proteins that are recalcitrant to recombinant expression *in vivo*, with several distinct advantages (Carlson, Gan, Hodgman, & Jewett, 2012). CFPS enables the synthesis of active protein in under a day, rapid screening of large protein libraries, and direct quantification of soluble and total yields of the target protein without cell lysis and purification. Moreover, the open reaction environment allows for precise control of reaction conditions, which is increasingly being applied to the biosynthesis of active membrane proteins. In recent years, a growing number of examples include the manufacture of ATP synthase (Matthies, Haberstock, Joos, Dotsch, & Al., 2011), G-protein-coupled receptors (Corin et al., 2011; Kaiser et al., 2008; Proverbio et al., 2013), and human mitochondrial voltage-dependent anion channel (Damiati et al., 2015). The key idea is to synthesize membrane proteins in the presence of natural or synthetic lipids and/or detergents that help solubilize the membrane protein. For example, membrane mimics such as micelles, liposomes, and nanodiscs have been used to synthesize membrane proteins in soluble, well-folded conformations (Cappuccio et al., 2008; Kubick, Gerrits, Merk, Stiege, & Erdmann, 2009; Liguori, Marques, & Lenormand, 2008; Matthies et al., 2011; Sachse, Dondapati, Fenz, Schmidt, & Kubick, 2014; Schwarz et al., 2007). Additionally, detergents such as *N*-dodecyl-β-D-maltoside (DDM) and Triton X-100 are also commonly used as additives to prevent aggregation of hydrophobic polypeptide sequences (Jaffee & Imperiali, 2013; Klammt et al., 2004; Lyukmanova et al., 2012; Seddon, Curnow, & Booth, 2004).

Given the emergence of efforts to synthesize membrane proteins with CFPS, we aimed to develop an *Escherichia coli* crude extract-based platform that combines CFPS and IVG for synthesis and characterization of OSTs (**Figure 1**). Although many platforms for CFPS have been used for membrane protein expression, including wheat germ (Periasamy et al., 2013), rabbit reticulocytes (Kaneda et al., 2009), and insect cells (Kubick et al., 2009; Sachse et al., 2013), the *E. coli* platform was chosen for three reasons. First, it has the highest batch protein biosynthesis yields, with up to 2.3 mg/mL reported for a model green fluorescent protein (Caschera & Noireaux, 2014). Second, *E. coli* lysates lack native *N*-linked glycosylation machinery, providing a blank canvas for bottom-up glycoengineering (Valderrama-Rincon et al., 2012) as well as eliminating the possible contamination from native glycosylation machinery. Third, emerging bacterial glycoengineering efforts have recently demonstrated the potential for using bacterial systems for fundamental and applied glycobiology efforts (for recent reviews see Baker, Çelik, & DeLisa, 2013 and Merritt, Ollis, Fisher, & DeLisa, 2013). The functional transfer of the *N*-linked protein glycosylation cluster from *Campylobacter jejuni* into *E. coli* set a precedent for engineering heterologous glycosylation pathways in *E. coli*, and has since enabled extensive study of protein glycosylation machinery for a variety of exciting applications (Baker et al., 2013; Valderrama-Rincon et al., 2012; Wacker et al., 2002). For example, PglB homologs from a variety of microbes have been expressed in *E. coli* and shown to glycosylate non-native target proteins with non-native LLOs *in vivo* (Ollis et al., 2015). Importantly for this work, *E. coli* has also served as a chassis for studying *in vitro* glycosylation of CFPS-expressed acceptor proteins with purified PglBs from recombinant expression cultures (Guarino & DeLisa, 2012).

**Figure 1:**
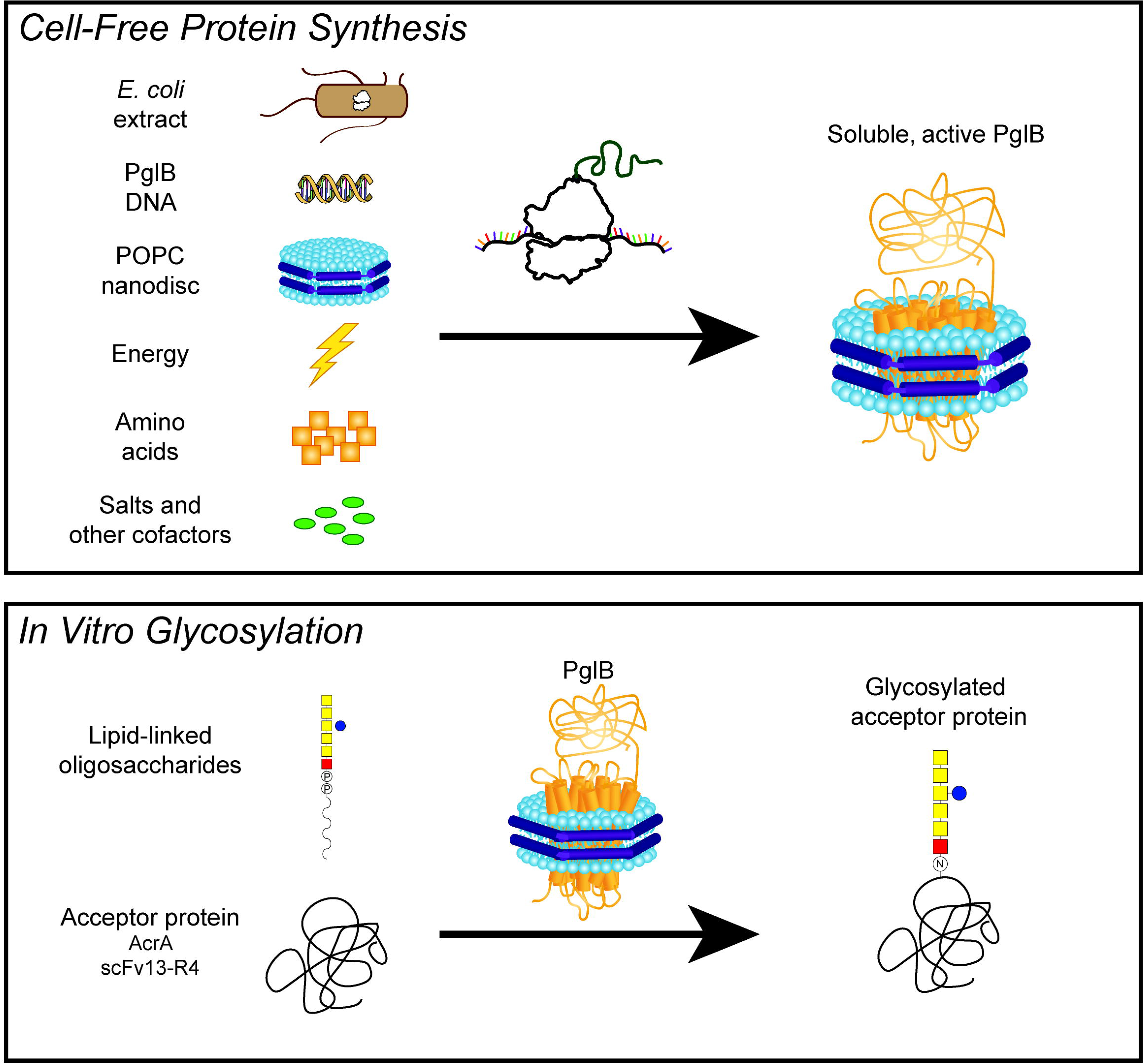
CFPS with membrane mimics allows expression of OSTs that are active *in vitro*. OSTs are first expressed in CFPS reactions containing membrane mimics. The synthesized OSTs can then be transferred directly to an IVG reaction containing LLOs and purified acceptor proteins. A key feature of this system is that LLOs and OSTs are derived from crude *E. coli* preparations, with no need for extensive purifications.

In this work, we extend our previous approach by demonstrating that the OST can be synthesized in CFPS, rather than purified from cells. Specifically, we optimized the utilization of detergents and membrane mimics to enable rapid and efficient cell-free expression of soluble PglB from *C. jejuni* (*Cj*PglB). By testing more than 15 independent conditions, we identified conditions capable of synthesizing *Cj*PglB at yields of up to 440 µg/mL, and showed that *Cj*PglB can be quantitatively added to *in vitro* activity assays with no purification or processing necessary. Using our optimized CFPS conditions, we additionally demonstrated the generalizability of our platform to the soluble synthesis of three additional bacterial OSTs, which actively glycosylated various target proteins and sequons. Our method provides flexibility for rapidly testing OST variants, manipulating physicochemical conditions, and decoupling glycosylation activity from cell toxicity constraints (Guarino & DeLisa, 2012). We expect that our approach will be useful for generating and assessing the activity of diverse OSTs through the use of well-defined experimental conditions.

## Results

### Effects of membrane mimics on solubility and yields of *Cj*PglB

The goal of this work was to develop a CFPS platform for the expression and characterization of OSTs. As a model OST, we chose to synthesize *Cj*PglB because its lipid-linked oligosaccharide and amino acid sequon specificities are well-studied (Chen et al., 2007; Ollis et al., 2015). Additionally, *Cj*PglB functions as a single subunit, making its study more tractable than multi-subunit OSTs found in higher organisms. The initial metric for success in comparing reaction conditions was high solubility of *Cj*PglB in CFPS. Total and soluble OST expression was measured by incorporating ^14^C-leucine during 15-μL combined transcription-translation reactions at 30°C. CFPS yields were measured before (total) and after (soluble) centrifugation, where the centrifugation step was used to remove insoluble product from the reaction mixture (**Figure 2**). To gauge the impact membrane mimics on *Cj*PglB solubility, we compared multiple conditions to a control reaction where no membrane mimics were present. In the control reaction, solubility of *Cj*PglB was just 11%, with 49 ± 5 µg/mL soluble and 442 ± 28 µg/mL total yields. This is shown in the first condition of **Figure 2A**.

**Figure 2:**
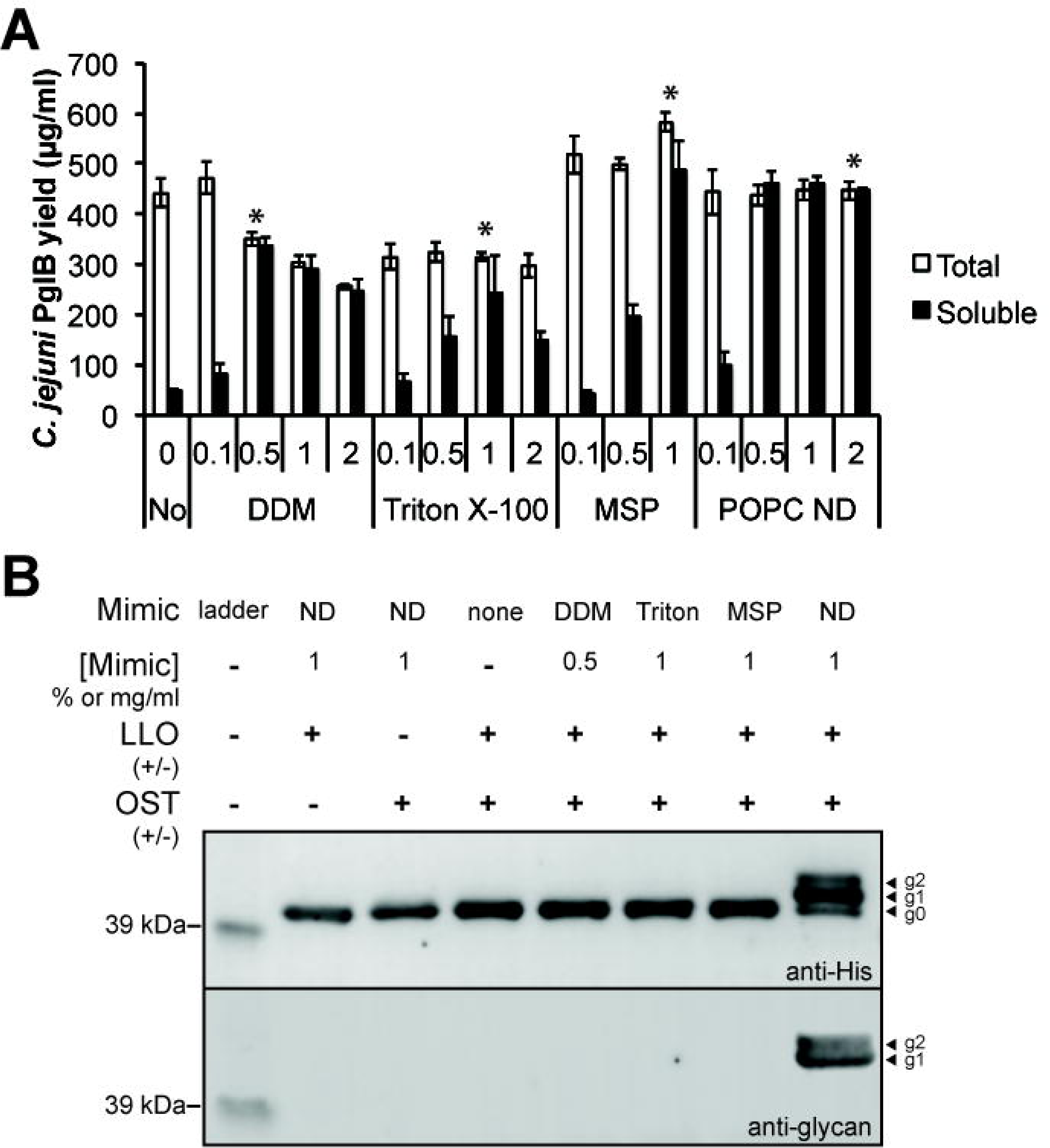
Soluble expression of active *Cj*PglB in CFPS reactions. (A) Total (white) and soluble (black) yields of PglB as measured by ^14^C-leucine incorporation supplemented with DDM, Triton X-100, MSP, and POPC nanodiscs (ND). 15-µL CFPS reactions were incubated at 30ºC for 6 h. Values represent the means and the error bars represent the standard deviations of three independent experiments. Asterisks indicate the conditions chosen for testing activity in an IVG reaction in (B). (B) CFPS-derived *Cj*PglB was tested for activity in IVG reactions at 30ºC overnight. *Cj*PglB at 0.65 µM was incubated with AcrA at 4.05 µM, and LLOs at 45% (v/v) from *E. coli* + pACYCpglB::kan membrane fractions. Blots were probed with anti-polyhistidine antibody and anti-glycan serum; activity is demonstrated by the appearance of singly (g1) and doubly (g2) glycosylated AcrA, which is confirmed by the anti-glycan blot. Full blots are shown in **Supplementary Figure 4**.

To increase soluble yields, CFPS reactions were supplemented with multiple membrane mimics at a range of concentrations. It is notable that these membrane mimics were supplemented at the start of the reaction to encourage proper folding of the hydrophobic domains within *Cj*PglB. The effects of adding non-ionic detergents to CFPS were studied first. DDM and Triton X-100 were chosen because they are commonly used in CFPS reactions (Klammt et al., 2004; Lyukmanova et al., 2012) and for extraction of membrane proteins from cellular membranes, particularly for functional assays (Jaffee & Imperiali, 2013; Moraes, Evans, Sanchez-Weatherby, Newstead, & Stewart, 2014; Seddon et al., 2004). Additionally, these detergents have routinely been added to IVG reactions with *Cj*PglB (Chen et al., 2007; Gerber et al., 2013; Guarino & DeLisa, 2012; Lizak et al., 2013; Slynko et al., 2009). With the exception of 0.1% (w/v) DDM, all conditions produced less total protein than was produced without a membrane mimic, but solubility was generally increased (**Figure 2A**). *Cj*PglB synthesized in the presence of DDM remained 95% soluble for 0.5, 1, and 2% (w/v) detergent conditions. The highest soluble yield of the detergent conditions tested was 338 ± 15 µg/mL for 0.5% (w/v) DDM and the best condition for Triton X-100, 1% (v/v), yielded 244 ± 71 µg/mL soluble *Cj*PglB at 77.6% solubility (**Figure 2A**).

We next tested the effect of nanodiscs on the synthesis of soluble *Cj*PglB in CFPS. Nanodiscs are defined nanostructures, composed of phospholipid bilayers that are solubilized into discoidal patches by two copies of an amphipathic membrane scaffold protein (MSP) (Bayburt & Sligar, 2010). Nanodiscs stabilize highly-hydrophobic protein domains by acting as a support for co-translational membrane association (Baumann et al., 2016; Bayburt & Sligar, 2010; Lyukmanova et al., 2012). Nanodiscs consisting of a 1-palmitoyl-2-oleoyl-sn-glycero-3-phosphocholine (POPC) lipid bilayer, encased by the MSP 1E3D1 were used. This combination led to the assembly of nanodiscs with a 12 nm average hydrodynamic diameter (**Supplementary Figure 1**) (Denisov, Grinkova, Lazarides, & Sligar, 2004). It is notable that 12 nm of lipid bilayer per nanodisc provides ample space to accommodate properly-folded PglB, which has a predicted diameter of roughly 5 nm across (Lizak et al., 2011; Musial-Siwek, Jaffee, & Imperiali, 2016). Mizrachi *et al*. have shown that fusing integral membrane proteins to MSP can be used to produce soluble, active membrane proteins (Mizrachi et al., 2015); hence, MSP in the absence of a lipid bilayer was also pursued as a potential hydrophobic shield. Both the MSPs and NDs allowed for total expression of at least 443 ± 44 µg/mL *Cj*PglB. Additionally, supplementing 1 mg/mL MSP to CFPS resulted in the product of 488 ± 58 µg/mL soluble *Cj*PglB; supplementing 0.5, 1, and 2 mg/mL NDs to reactions yielded 461 ± 22, 460 ± 15, and 447 ± 4 µg/mL soluble *Cj*PglB, respectively (**Figure 2A**). Further, the OST was shown to be associated with the nanodiscs through an immunoprecipitation assay (**Supplementary Figure 2**). Additionally, the time and temperature dependence of solubility was studied with ^14^ C-leucine incorporation to optimize the incubation duration and temperature of CFPS reactions (**Supplementary Figure 3**).

We next tested the effectiveness of each of the membrane mimics in producing functionally active OST. Reaction products corresponding to the highest soluble conditions for each membrane mimic were added to IVG reactions as an activity assay. IVG reactions consisted of CFPS-derived *Cj*PglB, purified AcrA (a native *C. jejuni* glycoprotein that contains two glycosylation acceptor sites), and a crude membrane extract from *E. coli* cells carrying the pgl pathway, which encodes a biosynthetic gene cluster to build LLOs bearing the *C. jejuni N*-linked glycan. The *C. jejuni* glycan is a heptasaccharide with the following structure: GalNAc-α1,4-GalNac-α1,4-[Glcβ1,3-]GalNAc-α1,4-GalNAc-α1,4-GalNac-α1,3-Bac-β1, where Bac is bacillosamine, or 2,4-diacetamido-2,4,6-trideoxyglucose (Young et al., 2002). OST activity was assayed via Western blots probed with an anti-polyhistidine antibody against the polyhistidine-tagged acceptor protein, or *C. jejuni* glycan-specific serum against the glycan. *Cj*PglB synthesized in CFPS reactions supplemented with nanodiscs exhibited glycosylation activity, as evidenced by bands in the anti-polyhistidine blot corresponding to singly glycosylated (G1) and doubly glycosylated (G2) forms of AcrA (**Figure 2B**; **Supplementary Figure 4**). Western blot analysis with anti-glycan serum further corroborated the attachment of the *C. jejuni* glycan to AcrA at either 1 or 2 sites. Specific glycosylation with the *C. jejuni* heptasaccharide at the two predicted sequons in AcrA was confirmed by liquid chromatography-tandem mass spectrometry (**Supplementary Figure 5**). Despite having greater than 95% soluble expression, *Cj*PglB produced by CFPS in the presence of 0.5% (w/v) DDM was completely inactive (**Figure 2B**; **Supplementary Figure 4**). Likewise, the absence of a membrane mimic, 1% (v/v) Triton X-100, and 1 mg/mL MSP also yielded inactive *Cj*PglB enzymes (**Figure 2B**). Importantly, no glycosylation was observed when LLOs were omitted from reaction mixture containing active *Cj*PglB (**Figure 2B**). Taken together, these data confirm that CFPS-derived *Cj*PglB produced in the presence of nanodiscs is active.

Having confirmed glycosylation activity, we took advantage of the open nature of CFPS and IVG reactions to optimize conditions for glycosylation efficiency and yields. By co-titrating crude LLO extract, purified acceptor protein, and CFPS reactions containing active *Cj*PglB, we quickly identified conditions that yielded high glycosylation efficiencies of another model glycoprotein called scFv13-R4^DQNAT^ (**Supplementary Figure 6**). The scFv13-R4^DQNAT^ protein is a single-chain Fv antibody containing a single optimized *C. jejuni* glycosylation sequon (DQNAT) behind a flexible linker as a C-terminal fusion (Fisher et al., 2011; Kowarik, Numao, et al., 2006; Silverman & Imperiali, 2016; Valderrama-Rincon et al., 2012). Under the described biochemical conditions, the highest conversion to glycosylated protein was observed when active *Cj*PglB was supplemented in 10-fold excess of purified scFv13-R4^DQNAT^ (*i.e.*, 2 µM OST and 0.2 µM purified acceptor protein).

### Synthesis of active bacterial OST homologs with CFPS

To test the generality of our platform, three additional bacterial OSTs, including PglB homologs from *Campylobacter coli* (*Cc*PglB), *Campylobacter lari* (*Cl*PglB) and *Desulfovibrio gigas* (*Dg*PglB), were expressed in CFPS containing 1 mg/mL nanodiscs using the optimized conditions from above. These OSTs have previously been demonstrated by Ollis *et al*. to glycosylate scFv13-R4^DQNAT^ with the *C. jejuni* glycan *in vivo* (Ollis et al., 2015). Each of these OSTs was produced with approximately 100% solubility as determined by ^14^ C-leucine incorporation (**Figure 3**) and the soluble yields were comparable to that achieved for *Cj*PglB. Specifically, we produced 439 ± 14 µg/mL or 5.3 ± 0.2 µM for *Cj*PglB, 326 ± 17 µg/mL or 4.0 ± 0.2 µM for *Cc*PglB, 339 ± 34 µg/mL or 4.0 ± 0.4 µM for *Cl*PglB, and 369 ± 32 µg/mL or 4.5 ± 0.4 µM for *Dg*PglB. Following soluble expression, we tested all four OSTs for their ability to glycosylate scFv13-R4^DQNAT^. Similarly to *Cj*PglB, all three OSTs were active (**Figure 4A**; **Supplementary Figure 7**), with *Cj*PglB and *Dg*PglB both having glycosylation efficiencies of >75% under the optimized IVG conditions tested.

**Figure 3:**
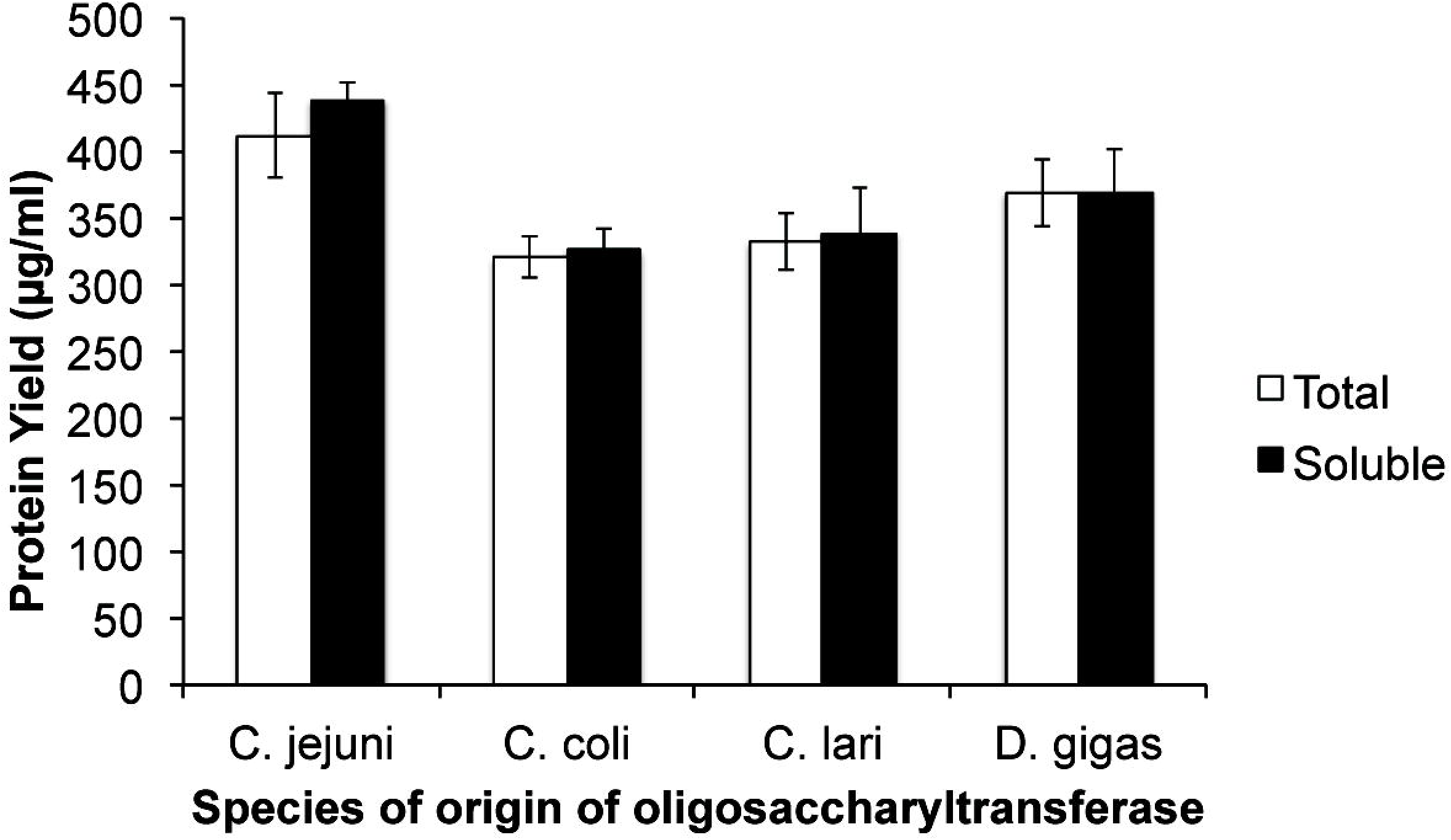
CFPS with nanodiscs yields soluble expression of four bacterial OSTs. Total (white) and soluble (black) yields of PglB homologs as measured by ^14^ C-leucine incorporation. 15-µl CFPS reactions were incubated at 30ºC for 6 h. Values represent the means and the error bars represent the standard deviations of six independent experiments.

**Figure 4:**
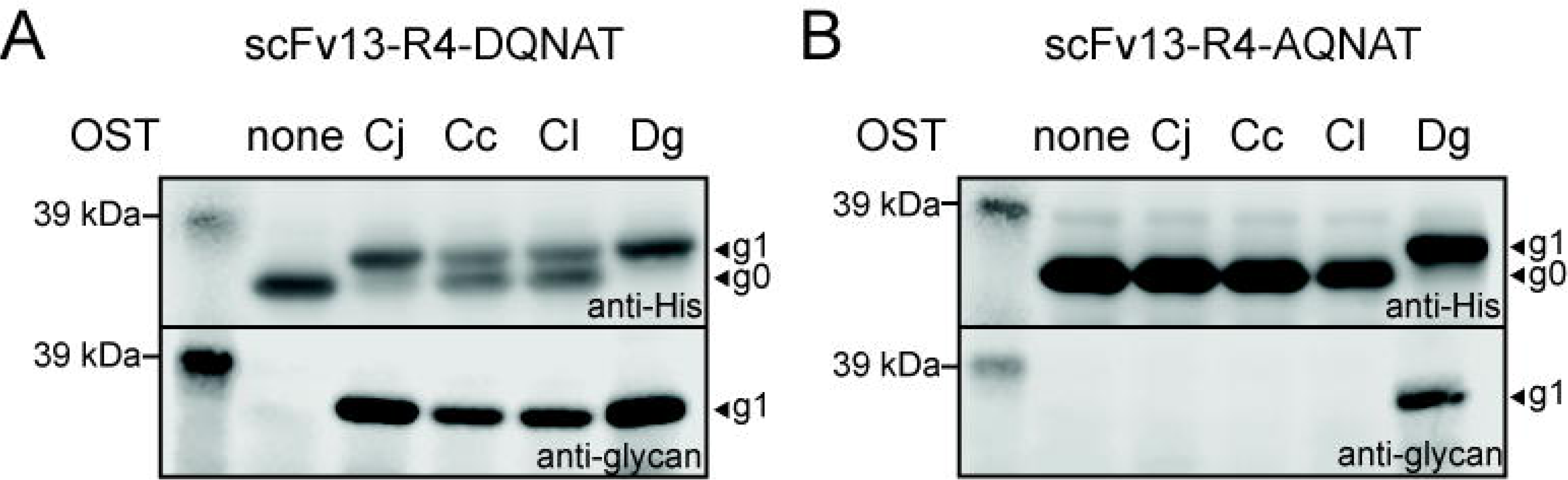
IVG of acceptor proteins with all four bacterial OSTs. CFPS-derived *Cj*PglB (Cj)*, Cc*PglB (Cc)*, Cl*PglB (Cl), and *Dg*PglB (Dg) were tested for activity in IVG reactions containing purified (A) scFv13-R4^DQNAT^ or (B) scFv13-R4^AQNAT^ as acceptor proteins. Blots were probed with anti-polyhistidine antibody and anti-glycan serum. Full blots are shown in **Supplementary Figures 7–8**.

We next set out to demonstrate how the open CFPS environment could facilitate rapid OST characterization. Specifically, we assessed sequon specificity of CFPS-derived OSTs by testing the glycosylation activity of the four bacterial OSTs with purified scFv13-R4^AQNAT^, which contains a single, C-terminal copy of the alternative glycosylation sequon, AQNAT (**Figure 4B**; **Supplementary Figure 8**). Because a key part of the catalytic cycle of the OST is to physically bind to the sequon before glycan transfer, simple modifications in the 5-amino acid sequon dictate the ability of the OST to exhibit glycosylation activity (*e.g.*, modification of DQNAT to AQNAT changes the -2 sequon position and eliminates activity of the *Cj*PglB on an otherwise identical protein) (Lizak et al., 2011; Ollis et al., 2015). Consistent with earlier *in vivo* experiments (Ollis et al., 2015), only CFPS-derived *Dg*PglB was able to glycosylate purified scFv13-R4^AQNAT^ *in vitro.* It is notable that when IVG reactions were run with CFPS reaction product containing no OST, model glycoproteins were not glycosylated (**Figure 4**). Taken together, our results demonstrate that protein glycosylation by CFPS-derived OSTs is robust, applicable to different proteins, and similar to activities displayed by their counterparts produced *in vivo.*

## Discussion

We present a new crude extract-based CFPS platform for the synthesis and characterization of OSTs. This platform was generated using an *E. coli*-based CFPS system augmented with nanodiscs to facilitate expression of four distinct, active bacterial OSTs at titers up to 440 µg/ml. A major benefit of the platform presented here is the ability to precisely control CFPS reaction conditions to obtain active PglB at high titers, offering an alternative to *in vivo* expression and purification for OST expression and study. By tuning CFPS conditions, we rapidly identified several conditions where greater than 75% soluble product was synthesized (**Figure 2A**). This open system allowed for identification of synthesis conditions that produced active and soluble products (**Figure 2B**) and the ability to test a variety of glycosylation conditions to drive higher glycosylation efficiencies. Finally, this method was extended to three additional OSTs that displayed high soluble expression and activity with no additional optimization of the reaction conditions.

Notably, solubility is not sufficient for enzymatic activity of OSTs, and activity is largely dependent on the ability of the membrane mimic to support proper folding of the transmembrane regions. These observations are supported by structural studies of bacterial and archaeal OST homologs (Lizak et al., 2011; Matsumoto, Taguchi, Shimada, Igura, & Kohda, 2017), which reveal that proper folding of transmembrane regions is crucial for activity. Specifically, several residues located between transmembrane regions are implicated in substrate recognition and the ability of an OST to act on the acceptor asparagine and lipid donor (Lizak et al., 2011). Since these residues must be precisely positioned with respect to the active site, proper folding of the hydrophobic regions is key for OST activity. OSTs produced co-translationally with detergents did not show glycosylation activity despite high solubility, and high yields of active enzyme were only achieved through CFPS supplemented with nanodiscs. We hypothesize that the phospholipid bilayer afforded by nanodiscs more closely mimics a native membrane environment than detergents do, providing the necessary lipid environment for activity. Generally, cell-free expression of membrane proteins using nanodiscs has been shown to lead to the direct incorporation of proteins into pre-formed nanodiscs, supporting our observations (Baumann et al., 2016; Denisov & Sligar, 2017; Lyukmanova et al., 2012). It is notable that PglB has been produced *in vivo* in *E. coli* and extracted with detergents (such as Triton X-100) while retaining activity. We hypothesize that *in vivo, E. coli* phospholipids support proper folding PglB, and remain associated with the enzyme through purification and detergent extraction, allowing the enzyme to remain in active conformation (Guarino & DeLisa, 2012; Jaffee & Imperiali, 2013). Ultimately, existing literature suggests that preferred membrane mimics vary between classes of membrane proteins and particular applications. Therefore, it is important for a synthesis platform to be compatible with a wide range of membrane mimics, as is the case in the current work.

In the future, we anticipate the use of our method for the synthesis and study of other single-subunit OSTs, especially given the speed, simplicity, and consistency of this CFPS method. An expression-based system for characterizing OSTs is needed, especially because of the significant challenges that exist in determining meaningful structure-function relationships (Maita et al., 2010). These challenges stem from the wide sequence divergence of OSTs. For example, although each of the homologs tested here are of proteobacterial origin, they differ in sequence homology from *C. jejuni* (82.7% sequence similarity for *C. coli*, 57.3% for *C. lari*, 16% for *D. gigas*) (Szymanski & Wren, 2005). Thus, the expression method developed here would be particularly valuable for characterization of OSTs which are poorly characterized or predicted by the deluge of recent microbial genomic sequencing efforts (Maita et al., 2010; Weerapana & Imperiali, 2006). Our method bypasses issues often encountered with *in vivo* OST characterization by decoupling enzyme yields from activity and avoiding protein purification. For instance, in work from Ollis, *et al*. several of the predicted bacterial OSTs analyzed in that study were deemed inactive (Ollis et al., 2015). However, it is difficult to determine whether this was due to poor *in vivo* expression, or due to poor affinity of the enzymes for the given substrates. The cell-free system developed here might be able to elucidate answers to such questions.

It is also plausible to extend a CFPS-based approach to high-throughput screening of OSTs for engineering novel functions and understanding their biology. Using evolution techniques, the *C. jejuni* PglB has already been engineered *in vivo* to accept the general eukaryotic N-X-S/T sequon as opposed to the more stringent native D/E-X_1_-N-X_2_-S/T sequon (Ollis, Zhang, Fisher, & DeLisa, 2014), to increase the enzyme’s efficiency (Ihssen et al., 2015), and to gain structural insight (Ihssen et al., 2012). However, glycan and lipid specificities are other useful attributes that could be tailored, given sufficient throughput of expression and analysis. With these targets, one could not only engineer novel OSTs, but also begin to probe which residues confer specific activities and specificities through experiments that would be a challenge to perform *in vivo* due to host cell lipid, and loss of library coverage *in vivo* (Ihssen et al., 2012). *In vitro* screening technologies could complement existing *in vivo* evolution methods, especially if integrated with liquid handling robotics for the generation and screening of enzyme variants with high-throughput.

In summary, CFPS systems are rapidly emerging as a powerful platform to understand, harness, and expand the powerful capabilities of biological systems. A recent surge of applications in prototyping genetic circuits (Noireaux, Bar-Ziv, & Libchaber, 2003; Takahashi et al., 2015), optimizing metabolic pathways (Dudley, Anderson, & Jewett, 2016; Karim & Jewett, 2016; Kay & Jewett, 2015), enabling portable diagnostics (Pardee et al., 2016), facilitating on-demand biomolecular manufacturing (Pardee et al., 2016; Salehi et al., 2016), and producing therapeutics at the commercial scale (Yin et al., 2012). Here, we show that CFPS can also be applied to the synthesis and characterization of OSTs. Poised at the intersection of glycobiology and synthetic biology, our platform has demonstrated rapid expression of a variety of soluble, active OSTs. These properties will allow for easy adoption of this platform to deepen our understanding of OST enzymology for advancing glycobiology.

## Materials and Methods

### Plasmid preparation

The open reading frames of PglBs and acceptor proteins were subcloned into the pET-based pJL1 vector by isothermal Gibson assembly. The Uniprot accession numbers for PglBs expressed in this study are Q9S4V7, A0A1B3X965, A0A0A8H643, and T2G1X6 for *C. jejuni, C. coli, C. lari*, and *D. gigas*, respectively. PglB genes were expressed with either a C-terminal FLAG-tag or an HA-tag. Acceptor proteins were tagged with a C-terminal polyhistidine tag for Western Blot analysis and purification. Plasmids were isolated for use in CFPS reactions using a maxi prep kit (Qiagen) followed by ethanol precipitation. Plasmids used for transformation were isolated for use via a miniprep kit (Zymo).

### Nanodisc preparation and characterization

Nanodiscs were prepared as described in Bayburt, *et al*. (Bayburt & Sligar, 2010). 1-palmitoyl-2-oleoyl-sn-glycero-3-phosphocholine (POPC) was purchased from Avanti Polar Lipids. Membrane scaffold protein 1E3D1 was purchased from Sigma-Aldrich. Nanodiscs were dialyzed into 1x PBS, and stored at -80ºC in aliquots. Nanodiscs were then analyzed on a Superdex 10/300 column at 0.5 mL/min in phosphate-buffered saline. The column was calibrated using an SEC calibration standard ranging over 15–600 kDa (Sigma).

### E. coli extract preparation

*E. coli* extracts were prepared as in Kwon & Jewett, 2015. BL21 (DE3) cells (Life Technologies) grown in 1 L of 2xYTPG media in full-baffle shake flasks at 37ºC. At an OD_600_ of 0.4, 1 mM of IPTG was added to induce T7 RNA polymerase production. Cells were harvested at an OD_600_ of 4.5. Cells were pelleted via centrifugation at 5000xg for 15 min at 4ºC, washed three times with cold S30 buffer (10 mM tris acetate, pH 8.2; 14 mM magnesium acetate; 60 mM potassium acetate; and 1 mM dithiothreitol), flash-frozen with liquid nitrogen, and stored at -80ºC. For lysis, cells were thawed on ice and resuspended in 1 mL of S30 buffer per gram cells, then lysed in an EmulsiFlex-B15 homogenizer (Avestin) in a single pass at a pressure of 22,500 psi. Cellular debris was removed by two rounds of centrifugation at 30,000xg for 30 min at 4ºC. The supernatant was incubated at 120 rpm for 80 min at 37ºC, then centrifuged at 15,000xg for 15 min at 4ºC. The final supernatant was flash-frozen with liquid nitrogen and stored at -80ºC until use. This extract contained 29.5 +/- 0.7 total protein as measured by a Quick-Start Bradford protein assay kit (Bio-Rad).

### Cell-free protein synthesis (CFPS)

CFPS reactions were performed with a modified, oxidizing PANOx-SP system (Jewett & Swartz, 2004; Zawada et al., 2011). 15 µL reactions were performed in 1.5 mL microcentrifuge tubes containing: 12 mM magnesium glutamate, 10 mM ammonium glutamate, 130 mM potassium glutamate, 1.2 mM adenosine triphosphate (ATP), 0.85 mM guanosine triphosphate (GTP), 0.85 mM uridine triphosphate (UTP), 0.85 mM cytidine triphosphate (CTP), 0.034 mg/mL folinic acid, 0.171 mg/mL E. coli tRNA (Roche), 2 mM each of 20 amino acids, 30 mM phosphoenolpyruvate (PEP, Roche), 0.33 mM nicotinamide adenine dinucleotide (NAD), 0.27 mM coenzyme-A (CoA), 4 mM oxalic acid, 1 mM putrescine, 1.5 mM spermidine, 57 mM HEPES, 13.3 µg/mL plasmid, and 30% (v/v) S30 extract. Membrane mimics were added as described above. These mimics included *n*-dodecyl-β-D-maltoside (DDM, Anatrace), Triton X-100, membrane scaffold protein 1E3D1, and 1-palmitoyl-2-oleoyl-sn-glycero-3-phosphocholine (POPC) nanodiscs. If not specified, the reactions contained 1 mg/mL POPC nanodiscs. Reactions were incubated for 6 h at 30ºC. Unless otherwise noted, reagents were purchased from Sigma-Aldrich. To quantify soluble and total protein yields, 0.50 µL (0.05 µCi) of radioactive ^14^ C-Leucine was added to each 15 µL CFPS reaction. After the reaction was complete, soluble proteins were separated by centrifugation at 20,000xg for 10 min at 4ºC. Proteins were quantified based on trichloroacetic acid (TCA)-precipitable radioactivity yields in a MicroBeta2 scintillation counter (PerkinElmer) (Calhoun & Swartz, 2005).

### Lipid-linked oligosaccharide crude membrane extract preparation

Plasmid pACYCpglB::kan (Wacker et al., 2002) bearing the glycosylation pathway for *C. jejuni* with the pglB gene inactivated by the insertion of a kanamycin resistance cassette was transformed into *E. coli* CLM24 cells and plated on selection plates (LB, 34 µg/mL chloramphenicol) for synthesis of *C. jejuni* LLOs (GalNAc-α1,4-GalNac-α1,4-[Glcβ1,3-]GalNAc-α1,4-GalNAc-α1,4-GalNac-α1,3-Bac-β1-UndPP). An overnight culture was started in 2xYTP media with 34 µg/mL chloramphenicol. One liter of 2xYTP (10 g/L yeast extract, 16 g/L tryptone, 5 g/L NaCl, 7 g/L potassium phosphate dibasic, 3 g/L potassium phosphate monobasic; pH 7.2) and antibiotic was inoculated at OD_600_ of 0.08. The strain expressed the glycosyltransferases (GTs) necessary for the synthesis of the *C. jejuni* lipid-linked oligosaccharide (LLO) on an undecaprenyl pyrophosphate anchor. Crude membrane extract was prepared as in Ollis et al., 2015. Briefly, cells were harvested by centrifugation, washed twice in resuspension buffer (50 mM Tris-HCl, 25 mM NaOH, pH 7.0), and finally resuspended in 10 mL of resuspension buffer per 1 g of cell pellet. The cell suspension was then frozen in liquid nitrogen and stored at -80ºC overnight. Cell pellets subsequently were thawed and lysed by homogenization with an EmulsiFlex-B15 homogenizer (Avestin). The resulting extract was centrifuged twice at 15,000xg for 20 min at 4ºC to remove unlysed cells. The supernatant was then ultra-centrifuged at 100,000xg for 1 h at 4ºC to pellet the lipid fraction. The pellet was resuspended in buffer with detergent (50 mM Tris-HCl, 25 mM NaCl, 1% (v/v) Triton X-100, pH 7.0) using a Dounce homogenizer. This suspension was incubated 1 h at room temperature, then centrifuged for 16,000xg for 1 h at 4ºC. The supernatant of this final spin was collected and stored at 4ºC for *in vitro* glycosylation reactions.

### Recombinant expression and purification of acceptor proteins

20 ng of plasmid DNA encoding acceptor proteins (scFv13-R4-D/AQNAT and AcrA) were transformed into BL21 Star (DE3) cells, plated on LB selection plates (50 µg/mL kanamycin), and grown overnight at 37ºC. Overnight cultures were inoculated from a single colony and grown overnight in 2xYTP with kanamycin. The following day, 100 mL 2xYTP with kanamycin was inoculated with saturated overnight cultures to a final OD_600_ of 0.08 and grown at 37ºC until OD_600_ of 0.6–0.8. Protein expression was induced with the addition of 0.5 mM IPTG and culture temperature was shifted to 30ºC. Cultures were incubated for 5 h before harvesting by centrifugation at 4,000xg for 15 min. Harvested pellets were frozen in liquid nitrogen and stored at -80ºC for further use.

After thawing pellets for 1 h on ice, pellets from 100 mL expression cultures were resuspended in 10 mL of wash buffer (50 mM sodium phosphate, 300 mM NaCl, pH 8) and collected at 4,000xg for 10 min. Pellets were washed twice before lysis. After the second wash step, cells were resuspended in 10 mL of lysis buffer (50 mM sodium phosphate, 300 mM NaCl, 10 mM imidazole, 1 mM phenylmethanesulfonyl fluoride, pH 8) and lysed via high-pressure homogenization with an EmulsiFlex-B15 homogenizer (Avestin). Resuspended culture was passed through the homogenizer for a total of 4 passes at 21,000 psi. Lysate was clarified by centrifugation at 15,000xg for 15 min at 4ºC. The supernatant of this centrifugation was removed, and centrifuged again at 15,000xg for 15 minutes. The supernatant of the second centrifugation was removed for further purification.

Polyhistidine-tagged acceptor proteins were purified from culture supernatants using Ni-NTA agarose resin or Ni-NTA spin columns (Qiagen) under manufacturer’s recommendations for native conditions. Resin or columns were washed with 3x packed resin volumes of wash buffer containing 20 mM imidazole, 50 mM imidazole, followed by elution with 500 mM imidazole. Eluted fractions were dialyzed into a storage buffer (50 mM HEPES, 500 mM NaCl, 1 mM EDTA, pH 7) and stored at 4ºC for subsequent use.

### In vitro glycosylation reactions

Ten microliter IVG reactions containing crude PglB expressed in CFPS, crude LLOs in *E. coli* membrane fraction, and purified acceptor protein were combined. CFPS reactions containing OSTs were diluted to 3.8 µM OST in CFPS pre-mix. 5 µL of this CFPS reaction was added to IVGs. 1 µL of purified acceptor protein at 2 µM in 500 mM NaCl, 30 mM Tris, and 1 mM EDTA, pH 7 were added to IVG reactions. 1 µL of a master mix (10% (w/v) Ficoll 400, 500 mM HEPES, pH 7.4, and 100 mM manganese chloride) was added just prior to incubation to reduce precipitation of reaction components. Reactions were then filled to volume with crude membrane extract containing LLOs. This gives the following concentrations of relevant components: 1.6 µM OST, 0.2 µM acceptor protein, 1% (w/v) Ficoll 400, 50 mM HEPES, and 10 mM manganese chloride. Reactions were incubated overnight at 30ºC.

### Western blot detection of *in vitro* glycosylation products

After incubation, IVG samples were centrifuged for five minutes at 20,000xg at 4ºC. Proteins in the supernatant were separated via SDS-PAGE on 4–12% Bis-Tris NuPAGE gels (Invitrogen) and transferred to Immobilon PVDF membranes (Millipore) with semi-dry transfer using a Trans-Blot SD (Bio-Rad) for Western blot analysis. A polyclonal, HRP-conjugated anti-polyhistidine antibody from rabbit (Abcam) was used to blot against acceptor proteins. The anti-glycan serum used was a generous gift from the Aebi Lab. WesternSure ECL reagents for chemiluminescence (LI-COR) or IRDye680 and IRDye800 secondary antibody conjugates (LI-COR) and imaged using Odyssey Fc imager (LI-COR).

### LC-MS/MS confirmation of presence and identity of glycosylation reaction products

AcrA samples were prepared for LC-MS/MS analysis by in-gel digestion as follows. Nickel column purified AcrA bearing two glycosylation sites was glycosylated in IVG reactions. Six IVG reactions were run in parallel as described above, then purified by Qiagen nickel affinity spin columns according to manufacturer specifications. After purification, the glycosylated AcrA was separated on a 4–12% SDS PAGE gel. The gel was then stained using InstantBlue (Expedeon). Gel slices containing both unglycosylated and glycosylated versions of AcrA were destained and subjected to in-gel trypsin digestion as previously described (Olsen, Mann, Shevchenko, Tomas, & Havlis, 2007) for at least 16 h, followed by extraction of tryptic peptides.

For confirmation of the presence of *C. jejuni* glycan on AcrA glycosylation sites, AcrA tryptic peptides were analyzed using an Oribtrap Fusion (Thermo Scientific) with Ultimate LC system equipped with a custom made C18 column and nanospray Flex Ion source. High-resolution MS scans and simultaneous detection of N-acetylhexosamine (HexNAc) fragment ions (+ 204 m/z) by MS/MS fragmentation confirmed the presence of *C. jejuni* glycans at both sites. For confirmation of the *C. jejuni* glycan identity, an identical sample of trypsin digested AcrA peptides was desalted using an Oasis HLB 96-well microelution plate (Waters Corp) according to manufacturer instructions. The eluent was then concentrated to a total volume of 8 μL using a Speedvac (Thermo Scientific). A 2 µL sample was analyzed by LC-MS/MS using an Agilent 1290 UPLC system equipped with an ACQUITY UPLC Peptide BEH C18 Column (300 Å pore size, 1.7 μm, 2.1 mm X 100 mm, Waters Corp) coupled to a QTRAP 6500 mass spectrometer (AB SCIEX). MS data acquisition and analysis was carried out using Analyst software (AB SCIEX). A precursor ion scan was carried out to identify, isolate, and fragment glycopeptides containing HexNAc residues similar to previous works (Zhang et al., 2012). Briefly, the 0.1% (v/v) formic acid (FA) in water and acetonitrile (ACN) were used as aqueous (A) and organic (B) solvents, respectively with a column temperature of 60°C. A 96 minute LC gradient from 2% to 50% ACN in 0.1% FA at 0.2 mL/min flow rate was used to separate glycopeptides. A precursor ion scan in positive ion mode monitoring the N-acetylhexosamine fragment ion (+ 204 m/z) using a step size of 0.2 Da was completed.

Based on an Enhanced Resolution (ER) MS scan, Information Dependent Acquisition (IDA) triggered an MS/MS Enhanced Product Ion (EPI) scan of the three highest intensity precursor ions between 400 and 2000 m/z with charges of +2 to +5 that produced the diagnostic HexNAc fragment ion. Rolling collisional energy was used for each MS/MS scan. Acquired MS and MS/MS spectra were inspected using custom MATLAB scripts for position and composition of glycan modifications.

## Acknowledgements

We thank Markus Aebi for the generous gift of the antiserum against the *C. jejuni* glycan. We acknowledge the Cornell University Biotechnology Resource Center (BRC) for data acquisition with their LC-MS/MS data acquisition. We thank the University of Illinois at Chicago and Chicago Biomedical Consortium for use of their LC-MS/MS instruments at the Mass spectrometry, Metabolomics & Proteomics Facility. We thank Kyle Wilcox for training in the preparation of nanodiscs. JAS was supported by the National Science Foundation Graduate Research Fellowship, grant number DGE-1324585. This work was also supported by the Defense Threat Reduction Agency (GRANT11631647), the David and Lucile Packard Foundation, the Chicago Biomedical Consortium with support from the Searle Funds at the Chicago Community Trust, the Dreyfus Teacher-Scholar program, and the National Science Foundation (MCB 1413563). The authors declare that they have no conflict of interest.

## Author Contributions

JAS and JMH designed, conducted, and analyzed OST synthesis and activity experiments. JCS, TJ, and AN designed and optimized experimental conditions. WK and JEK designed, performed, and interpreted mass spectrometry experiments. MCJ and MPD directed the studies and interpreted data. JAS, JMH, and MCJ wrote the paper with assistance from JCS, WK, JEK, TJ, AN, and MPD.

## Abbreviations

CFPS – cell-free protein synthesis, IVG – *in vitro* glycosylation, LLO – lipid linked oligosaccharide, OD – optical density, OST – oligosaccharyltransferase, scFv – single chain variable fragment, ND – nanodiscs

